# Modeling MEK inhibitor-Associated Retinopathy *in vitro* using human induced pluripotent stem cell-derived retinal pigment epithelial cells

**DOI:** 10.1101/2025.10.03.680301

**Authors:** Lola P. Lozano, Madeleine Jennisch, Renato Jensen, Jason A. Ratcliff, Luke A. Wiley, Brynnon Harman, Megan J. Riker, Allison T. Wright, Bradley A. Erickson, H. Culver Boldt, Timothy M. Boyce, Robert F. Mullins, Elaine M. Binkley, Budd A. Tucker

## Abstract

Pharmacologic inhibitors of MEK are important anti-cancer drugs but can result in MEK inhibitor-Associated Retinopathy (MEKAR) in which vision is lost due to serous retinal detachments that form via an unknown mechanism. We hypothesized that the cause of this side effect is drug-induced dysfunction of retinal pigment epithelial (RPE) cells. To test this hypothesis, we used human induced pluripotent stem cell-derived RPE cells. We treated mature, hiPSC-derived RPE cells with selumetinib and measured impacts on RPE-specific function, structure, and gene expression. Selumetinib increases the ability of hiPSC-derived RPE to internalize bovine rod outer segments (1.9 vs 3.0, p=0.0024). It also decreases expression of aquaporin 1 during the first 10 days of treatment (2.7 vs 1.1, p=0.0015). It has no effect on the ability of hiPSC-derived RPE to maintain membrane integrity. Selumetinib alters gene expression of hiPSC-derived RPE, with significant changes in genes involved in transport of ions and small molecules regulating cell volume and lysosomal acidification. Selumetinib may lead to subretinal fluid accumulation by both increasing secretions into this space and decreasing outflow.

## INTRODUCTION

Many cancers develop activating mutations in the cell signaling mitogen-activated protein kinase/extracellular signal-regulated kinase (MAPK/ERK) pathway which lead to uncontrolled cell survival and proliferation (**Supplementary Figure 1A**) (1). Mutations in *MEK* frequently occur in cutaneous melanoma, non-small cell lung cancer, and pancreatic, colorectal, breast, and liver cancers. Four MEK inhibitors have received FDA approval for clinical use: cobimetinib for malignant melanoma with BRAF^V600^ mutations, binimetinib and trametinib for metastatic melanoma with BRAF^V600E/K^ mutations, and selumetinib for patients with Neurofibromatosis Type 1 and stage III or IV differentiated thyroid cancer. There is also evidence suggesting the efficacy of MEK inhibitors in the treatment of the basal subtype of bladder cancer and metastatic castration resistant prostate cancer (2, 3).

Unfortunately, due to the ubiquity of the MAPK/ERK pathway, MEK inhibitor therapy has a substantial number of side effects, some of which are quite significant. For instance, one of the more common side effects reported, affecting up to 90% of patients (4), is MEK inhibitor-Associated Retinopathy (MEKAR). On average, MEKAR manifests within the first two weeks of starting MEK inhibitor therapy (5). The retinopathy is typically multifocal and bilateral with findings isolated to the posterior pole along the main vascular arcades and macula. There is often at least one focus localized to the fovea, which causes loss of high acuity, central vision (5). In all cases of documented MEKAR and retinopathies due to other drugs inhibiting the MAPK/ERK pathway, fluid accumulates between the ends of the photoreceptor outer segments at the interdigitation zone with an intact retinal pigment epithelium (RPE) (**Supplementary Figure 1B-C**) (5–7). This fluid accumulation is not due to abnormal fluid leakage from the choroidal vasculature, as seen in central serous chorioretinopathy, which is demonstrated by no leakage of dye on fluorescein angiography (4, 5, 8–10). While MEKAR often resolves spontaneously by approximately 32 days after starting treatment, a subset of patients have persistent retinopathy and severe visual side effects requiring their medication be discontinued (5, 10).

Although the mechanism of MEKAR is unknown, given the robust clinical phenotype (subretinal fluid accumulation without leakage), it is postulated to be the result of drug-induced RPE cell dysfunction. RPE cells, which form a monolayer below the neural retina above the choroidal vasculature, are critical for maintaining retinal homeostasis (**Supplementary Figure 1D-E**). For instance, RPE cells play a key role in phagocytosing shed photoreceptor outer segments (POS), scavenging reactive oxygen species secreting growth factors for both the retina and underlying choroid, and contributing to the choroid-retinal barrier via tight junction proteins (zonula occludens, occludins, claudins) and transporting molecules between these two structures (11–13). Their role in molecular trafficking, maintaining subretinal fluid volume as well as the proper ionic concentrations for photoreceptors to function, exists because of the unique protein polarity on the basolateral versus apical side of the RPE plasma membrane (12). We hypothesize that drug-induced disruption of these unique RPE cell functions causes the pathologic subretinal fluid accumulation observed in patients with MEKAR.

To investigate, we generated human induced pluripotent stem cell-derived RPE cells and treated them with the clinically used MEK inhibitor, selumetinib. As most patients manifest MEKAR within the first two weeks of beginning MEK inhibitor therapy, and many individuals experience resolution by 1-month, we collected data after treating cells for 10, 20, and 30 days. While we did not see an appreciable reduction in barrier function as determined by transepithelial resistance, by 10 days of treatment we did detect a reduction in aquaporin expression, increased photoreceptor cell outer segment phagocytosis, and disruption of transcriptional profiles associated with fluid flux and ion transport.

## RESULTS

### Generation of hiPSC-derived RPE cells

Donor fibroblasts were reprogrammed to human induced pluripotent stem cells (hiPSCs) via Sendai virus mediated expression of the transcription factors OCT4, SOX2, KLF4 and C-Myc as previously described (14). At 25-30 days following Sendai viral transduction hiPSC colonies were manually isolated and clonally expanded in Essential 8 media on laminin-521 coated cell culture surfaces. At passage 10 karyotypic analysis was performed to ensure lack of chromosomal abnormalities (**Figure 1A**).

**Figure 1.**
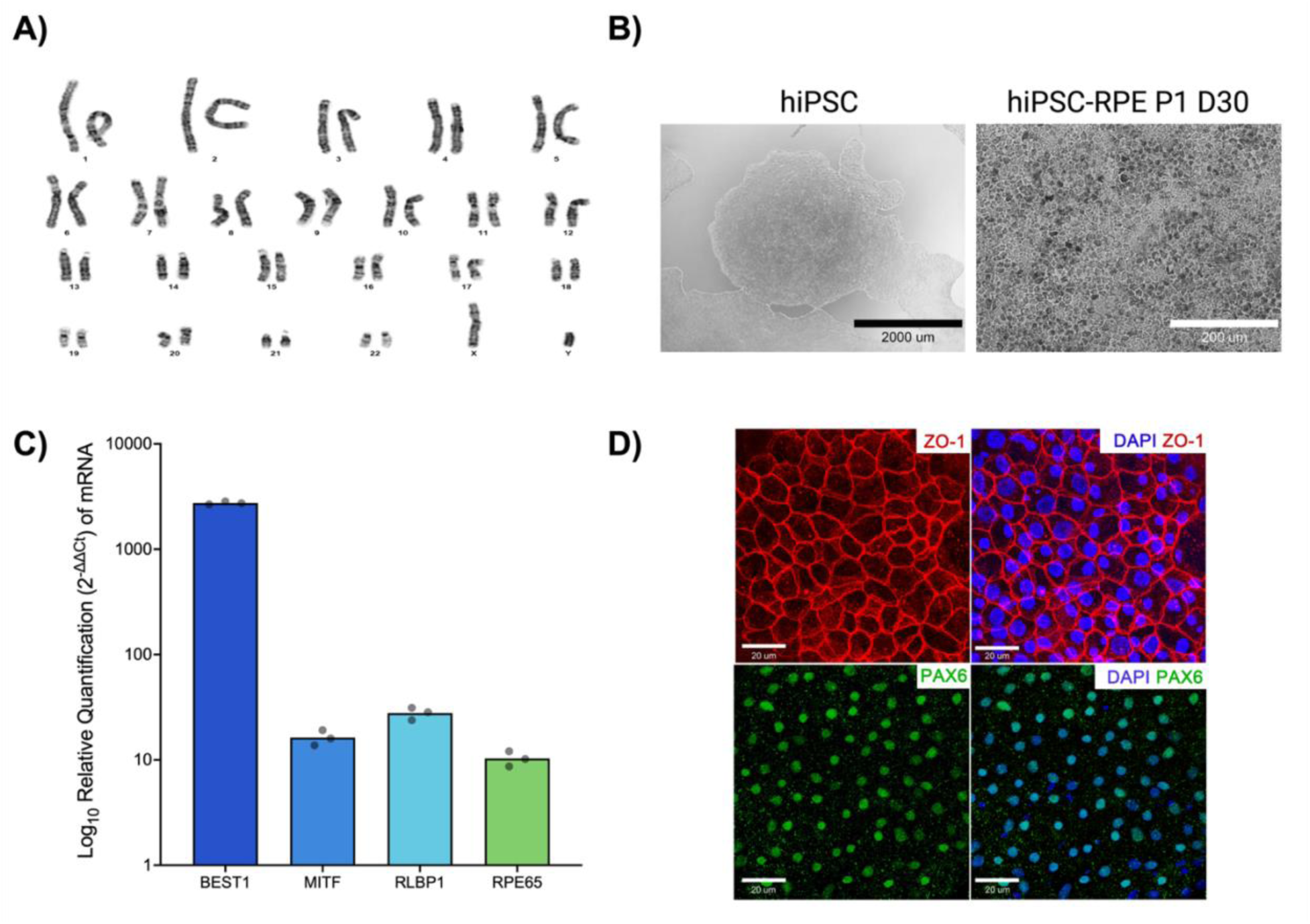
Differentiation of hiPSCs into RPE. **A)** Normal karyotype after reprogramming donor fibroblasts into human induced pluripotent stem cells (hiPSCs). **B)** hiPSCs display typical translucent colonies composed of densely packed cells. After differentiation to hiPSC-derived RPE, the cells form a cobblestone-like monolayer of pigmented polygonal cells, morphology characteristic of RPE cells. **C)** RT-qPCR for signature RPE genes. delta-delta Ct values with bar height at the mean of technical triplicates. Values are normalized to the housekeeping gene *B2M* and values from hiPSCs derived from the same donor. **D)** Immunocytochemistry for signature RPE proteins. Cell membranes label strongly for the tight junction protein, ZO-1, while nuclei show robust labeling for PAX6, a key protein in eye specific development.

Following confirmation of normal karyotype, hiPSCs were differentiated into retinal pigment epithelia (RPE) cells using the differentiation protocol developed by Foltz *et. al*. (15). By 30 days of differentiation hiPSC-derived RPE cells formed a monolayer of pigmented, polygonal cells with cobblestone morphology (**Figure 1B**). Expression of the RPE genes *BEST1*, *MITF*, *RLBP1*, and *RPE65* were confirmed using RT-qPCR (**Figure 1C**). In addition, expression of the tight junction marker ZO-1 and transcription factor PAX6 were confirmed via immunocytochemistry (**Figure 1D**).

### 10 μM selumetinib optimally inhibits phosphorylation of MAPK in hiPSC-derived RPE

To identify the dose of selumetinib suitable for suppressing MEK signaling without inducing widespread cell death, an MTS cell viability assay was performed on the hiPSC-RPE cells. In this experiment a log10-fold dilution of selumetinib ranging from 0.001 – 1000 μM was evaluated. hiPSC-RPE were treated for 72 hours before absorbance at 490 nm was determined by a microplate reader. As shown in **Supplementary Figure 2A**, cell viability dropped precipitously when selumetinib concentrations exceeded 100 μM. To determine if selumetinib doses of 1 μM and 10 μM (1 and 2 log units below the viability threshold) were sufficient to inhibit MEK signaling, Western blot analysis using antibodies targeting the MEK effector MAPK (**Supplementary Figure 2A**) was performed. While MAPK phosphorylation was reduced in cells treated with 1 μM selumetinib, near complete inhibition of MAPK phosphorylation was achieved using 10 μM selumetinib (**Supplementary Figure 2B**). To determine the impact of selumetinib on MAPK phosphorylation as a function of treatment time, hiPSC-derived RPE cells were evaluated following 10, 20, and 30 days of treatment. Relative to untreated controls, hiPSC-derived RPE cells cultured in media containing 10 μM selumetinib displayed significantly decreased MAPK phosphorylation at all time points evaluated: Days 10 – 30 combined, 0.86 ± 0.083 vs 0.20 ± 0.206 (**Supplementary Figure 2C**); Day 10, 0.66 ± 0.11 vs 0.15 ± 0.033 (**Supplementary Figure 2D**); Day 20, 0.71 ± 0.092 vs 0.15 ± 0.028 (**Supplementary Figure 2E**); and Day 30, 1.2 ± 0.17 vs 0.31 ± 0.056 (**Supplementary Figure 2F**).

### Selumetinib does not affect transepithelial electrical resistance of hiPSC-derived RPE cells

If selumetinib disrupts the ability of RPE to maintain a monolayer of cells connected via tight junctions, this could lead to non-selective movement of water and ions across the blood-retinal barrier. The integrity of this barrier can be tested by measuring the transepithelial electrical resistance (TEER) across a monolayer of RPE. Thus, to evaluate TEER in our system, hiPSC-derived RPE were plated onto transwell inserts. Beginning five days after plating (Day 0 on **Supplementary Figure 3**), cell media was changed following treatment condition and TEER measurements were recorded every 3 days for up to 12 days with a final measurement at day 19 using the EVOM Meter following the manufacturer’s protocol. No significant difference in mean transepithelial resistance was detected between untreated and selumetinib-treated hiPSC-derived RPE cells at any of the time points evaluated (**Supplementary Figure 3**).

### Selumetinib treatment increases internalization of bovine rod outer segments by hiPSC-derived RPE cells

Another critical function RPE carry out to maintain retina homeostasis is phagocytosis of photoreceptor outer segments. Diseases with disruptions to this process cause blindness (16, 17). To determine if selumetinib treatment impacts the ability of RPE to phagocytose photoreceptor outer segments, mature hiPSC-derived RPE cells cultured in the presence or absence of 10 μM selumetinib were incubated for 3 hours with media containing bovine rod outer segments (10 bROS/cell). Cells were subsequently washed and collected as described below. The process of outer segment phagocytosis by the RPE involves outer segment binding followed by internalization and lysosomal degradation. The phagocytosis conditions that were collected for analysis in this experiment included Total bROS (combined bound and internalized factions), Internalized bROS (the portion of bROS inside the cell), and 3- and 24-hour Chase conditions, which measure the amount of bROS present after allowing an additional 3- or 24-hours of culture time following bROS washout (i.e. to allow for degradation). To quantify bROS levels, western blot analysis for the rod outer segment protein rhodopsin (RHO) was used (**Figure 2A**). Compared to untreated controls, selumetinib-treated hiPSC-derived RPE cells were found to have significantly increased levels of internalized RHO (**Figure 2B**, 1.879 ± 0.967 vs 2.972 ± 1.028, p=0.024). When segregated by the number of days cells received treatment, we found the most consistent increase was detected in cells treated for the first 10 days (**Figures 2C**, 0.7 ± 0.14 vs 1.6 ± 0.25, p=0.021). While not reaching statistical significance, increased RHO abundance was still apparent following 20 days of treatment (**Figure 2D**, 1.3 ± 0.28 vs 2.7 ± 1.0, p=0.238). In the Day 30 treatment group, internalized RHO expression was significantly higher in the selumetinib-treated cells (**Figure 2E**, 3.3 ± 3.0 vs 4.4 ± 3.1, p=0.014). There was no statistically significant difference between treatment groups in any of the other phagocytosis conditions evaluated at any individual time point or all time points combined (**Figure 2C-E**). Collectively, these data demonstrate that treatment of hiPSC-derived RPE cells with 10 μM selumetinib enhanced rod outer segment phagocytosis but did not appear to alter clearance capacity (i.e., by just 3 hours following washout the levels of RHO inside the cells was not significantly different between selumetinib treatment and control conditions).

**Figure 2.**
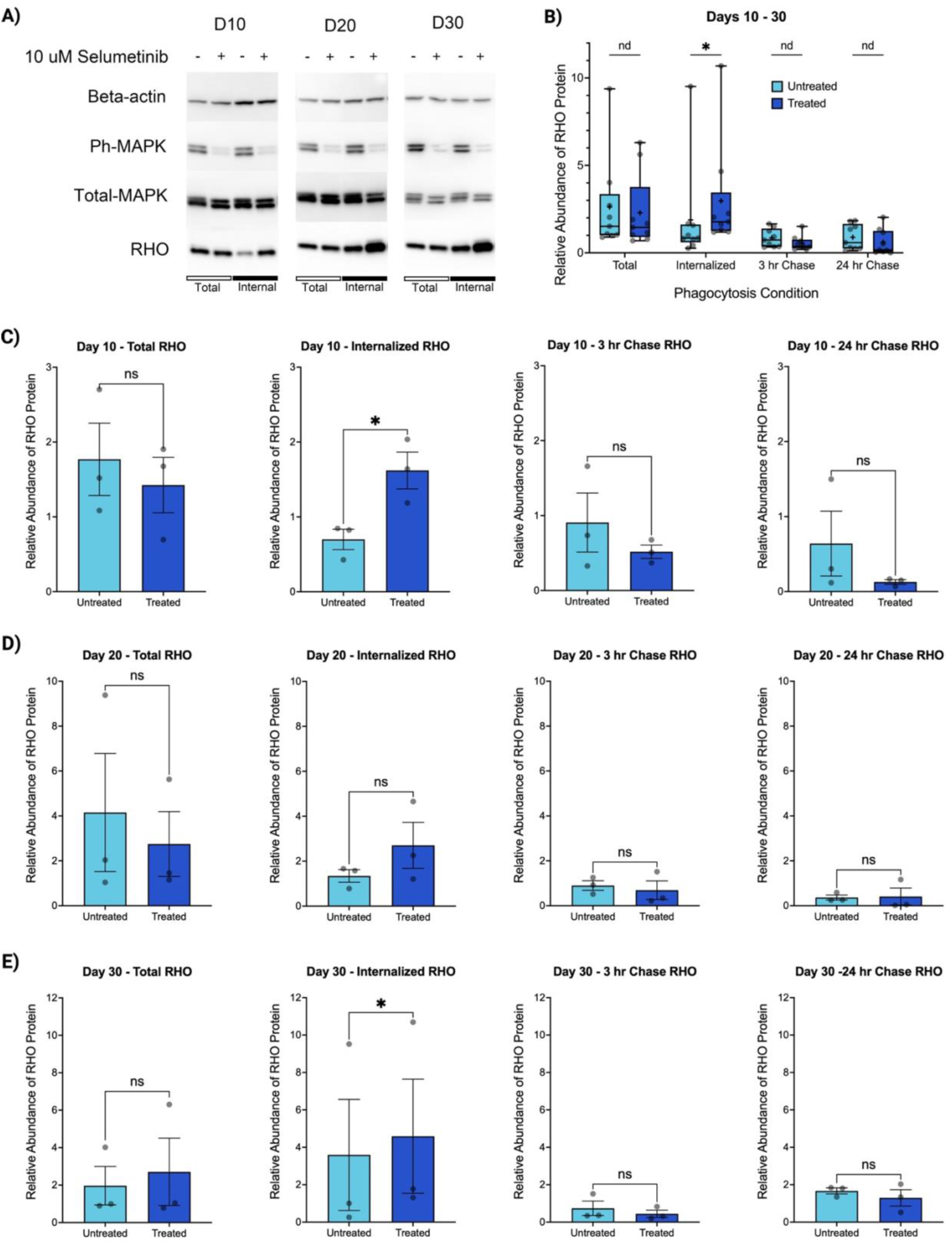
Selumetinib treatment increases bROS internalization by hiPSC-derived RPE cells. **A)** Representative Western Blot of Total and Internalized phagocytosis conditions show an increase in internalized RHO in hiPSC-derived RPE treated with 10 μM selumetinib versus untreated controls. **B)** When all treatment times were combined, 10 μM selumetinib significantly increases the internalization of RHO. There was no significant difference between treatment conditions for other phagocytosis conditions measured. Multiple paired t tests, Mean ± SEM, n = 9 pairs per phagocytosis condition. **C)** At Day 10, 10 μM selumetinib significantly increases the internalization of RHO and does not affect other phagocytosis conditions. **D)** At Day 20, there is no significant difference in relative abundance of RHO between treatment groups given phagocytosis conditions. **E)** At Day 30, 10 uM selumetinib significantly increases the internalization of RHO and does not affect other phagocytosis conditions. **C-E)** Results of paired t tests, Mean ± SEM, n = 3 pairs for each phagocytosis condition from independent experiments. *, p<0.05. Abbreviations: D10, 10 days of selumetinib treatment; D20, 20 days of selumetinib treatment; D30, 30 days of selumetinib treatmenr; nd, no discovery; ns, not significant.

### Selumetinib treatment decreases Aquaporin 1 (A*PQ1*) expression in hiPSC-derived RPE cells

AQP1 is principally expressed apically in RPE cells and helps prevent and decrease accumulation of subretinal fluid (18, 19). It has been shown to be regulated by the MAPK pathway in these cells (20). Because of this, AQP1 is hypothesized to play a role in MEKAR pathophysiology (5). To quantify changes in AQP1 protein expression between untreated hiPSC-derived RPE and those treated for 10-, 20-, and 30-days, western blot analysis was performed. When all treatment times were combined, we detected a statistically significant decrease in AQP1 levels in the selumetinib-treated hiPSC-derived RPE cells compared to untreated controls (**Figure 3A**, 1.6 ± 0.44 vs 0.98 ± 0.32, p=0.044). When the data was segregated by treatment time, it became evident that the impact on selumetinib on AQP1 expression occurred within the first 10 days of selumetinib treatment (**Figure 3B**, 2.7 ± 0.12 vs 1.1 ± 0.17, p=0.0015). Specifically, no significant difference in AQP1 expression was found following 20- (**Figure 3C**, 1.6 ± 1.1 vs 1.4 ± 0.95, p=0.252) or 30-days of selumetinib treatment (**Figure 3D**, 0.46 ± 0.039 vs 0.43 ± 0.069, p=0.406).

**Figure 3.**
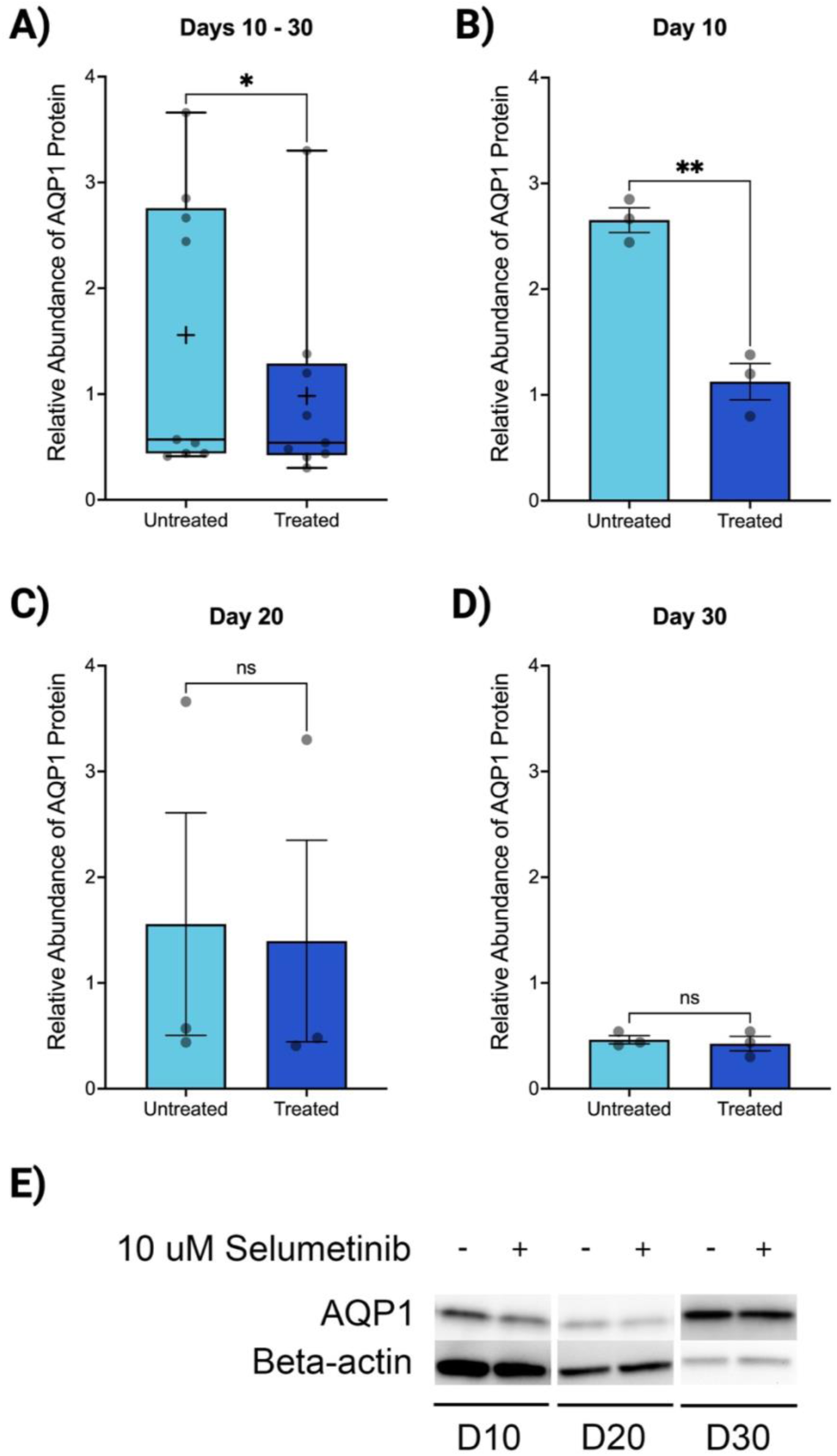
Selumetinib decreases aquaporin 1 (AQP1) expression. **A)** AQP1 expression in hiPSC-derived RPE cells in combined 10- (B), 20- (C) and 30-day selumetinib treatment groups versus untreated controls. For this analysis all treatment times were combined. **B-D)** AQP1 expression in hiPSC-derived RPE cells following 10- (B), 20- (C) and 30-days of selumetinib treatment versus untreated controls. Bars represent Mean ± SEM, n = 3 pairs from independent experiments for each time point. *, p<0.05; **, p<0.002. **E)** Representative Western blots depicting decreased AQP1 protein levels in hiPSC-derived RPE cells in selumetinib treated versus untreated controls following 10-, 20-, and 30-days of treatment. AQP1 was significantly decreased following 10 days of selumetinib treatment. The levels of AQP1 rebounded to normal by 20- and 30-days of selumetinib treatment.

### Selumetinib alters transcriptome of hiPSC-derived RPE

Because our assays measured specific functions and proteins we hypothesized could be affected by MEKAR pathophysiology, we decided to evaluate drug-induced changes to the hiPSC-RPE transcriptome. This analysis allowed us to analyze the pathophysiology in an unbiased nature and generate additional hypotheses regarding the mechanism of MEKAR. Bulk RNA-sequencing revealed significant changes in differentially expressed genes (DEGs) between untreated and selumetinib-treated hiPSC-derived RPE cells (**Figure 4A-B**). Of the 9,360 genes with measured expression, 957 genes were differentially expressed between the treatment and no treatment control groups. Notably, there was a significant decrease in all the top 50 DEGs associated with small molecule and ion transport and lysosomal acidification (*LRRC8C, CLCN5, SLC16A1, SLC6A6, SLC22A23, SL12A2, ATP6V1C2*) in selumetinib-treated hiPSC-derived RPE (**Figure 4A-B, Supplementary Table 1**). Ingenuity pathway analysis revealed the most significantly altered pathways were associated with neuroactive ligand-receptor interactions, calcium signaling, PI3K-Akt signaling, and extracellular matrix-receptor interaction (**Figure 4C**). Gene Ontology analysis revealed the top biological processes altered between treatment conditions were associated with visual perception, transport across the blood-brain barrier, and positive regulation of cytosolic calcium ion concentration, peptidyl-tyrosine phosphorylation, and ERK1 and ERK2 cascade (**Figure 4D**). Treatment with selumetinib significantly altered gene expression of several aquaporins: *AQP3, AQP4,* and *AQP7* increased and *AQP1* and *AQP11* decreased in cells treated with selumetinib (**Supplementary Table 2**).

**Figure 4.**
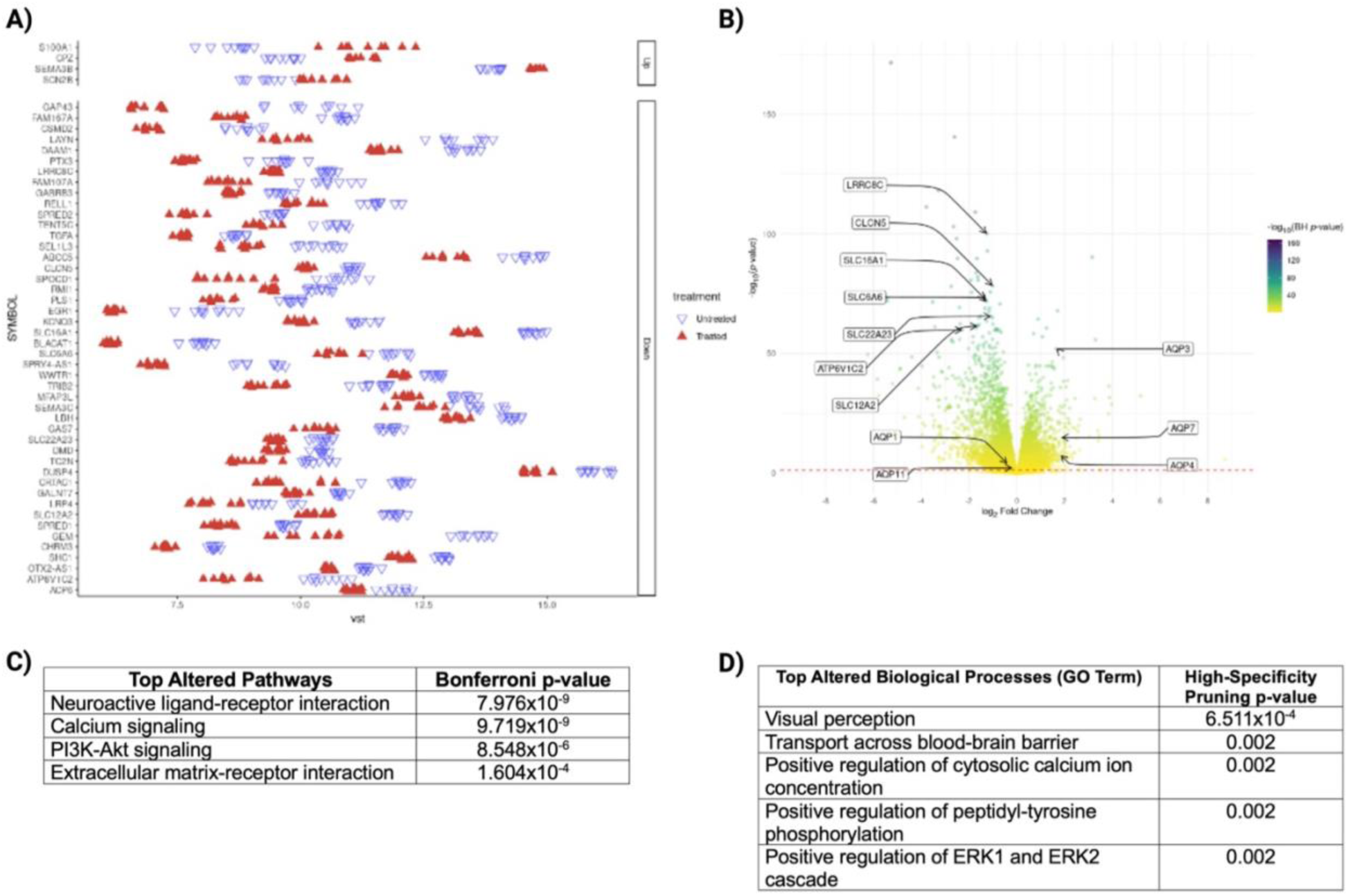
Selumetinib significantly alters gene expression in hiPSC-derived RPE cells. **A)** Beeswarm plot of top 50 DEGs. Y-axis ordering is determined by Benjamini-Hochberg (BH) adjusted p-value, with the lowest p-value at the top. **B)** Volcano plot of DEGs. Negative log_10_ of BH adjusted p-values. Estimates of fold change and significance were calculated from a pairwise contrast of treatment groups. Dashed red line indicates a p-value threshold of 0.05. **C)** Summary of top pathways altered genes between treatment groups using iPathwayGuide scores. **D)** Summary of top biological processes altered between treatment groups using Gene Ontology (GO) Analysis. Abbreviations: VST, variance stabilizing transformation; DEG, differentially expressed genes; GO, gene ontology; D10, 10 days of selumetinib treatment; D20, 20 days of selumetinib treatment; D30, 30 days of selumetinib treatment.

### Selumetinib does not alter hiPSC-derived RPE cell structure

We imaged cultured cells to determine if selumetinib affects hiPSC-derived RPE cell structure. Scanning electron microscopy (SEM) and transmission electron microscopy (TEM) visualized the external and internal compartments of the cells, respectively. No difference in cell structure was evident between untreated and selumetinib-treated hiPSC-derived RPE cells (**Supplementary Figure 4**).

## DISCUSSION

In the present study, we aimed to assess the effect of a commonly used MEK inhibitor, selumetinib, on hiPSC-derived RPE cells *in vitro*. We found that treatment of hiPSC-derived RPE with 10 μM selumetinib resulted in increased levels of internalized bovine rod outer segments (bROS) and decreased aquaporin-1 (AQP1) protein expression compared to untreated controls. These effects were most pronounced following 10 days of treatment and appeared to decrease as treatment times increased, which is consistent with clinical findings in patients with resolving MEKAR. Bulk RNA-sequencing revealed significant changes in gene expression. Notably, there was a significant decrease in expression of genes involved in transport of ions and small molecules regulating cell volume and lysosomal acidification. We did not find an effect of selumetinib on cell structure or the ability of hiPSC-derived RPE to maintain barrier function as determined by transepithelial electrical resistance measurements.

### Selumetinib increases internalization of bovine rod outer segments by hiPSC-derived RPE cells

In a daily cycle, each rod photoreceptor cell sheds 10% of its membranous discs contained within the photoreceptor outer segment. Shed outer segments are phagocytosed by RPE cells where they are converted into fatty acids and ketone bodies (13). Ketone bodies, primarily β-hydroxybutyrate (β-HB), are recycled back to the subretinal space for use in metabolism by photoreceptor cells. In this study we found that treatment of hiPSC-derived RPE cells with 10 μM selumetinib increased rod outer segment phagocytosis. We predict that increased internalization and degradation of photoreceptor outer segments leads to increased production and secretion of ketone bodies into the subretinal space *in vivo*. This would be expected to contribute to subretinal fluid accumulation characteristic of MEKAR. First, ketone anions attract sodium and water (21). Second, β-HB has been shown to increase vascular permeability via increased vascular endothelial growth factor (VEGF), which is the main factor that can increases blood-retinal barrier permeability (22, 23). However, this second mechanism is less likely to contribute to subretinal fluid accumulation in MEKAR as vascular permeability has not been detected with fluorescein angiography (4, 5, 8–10). Future work is needed to determine if MEK inhibitors increase ketone secretion by RPE cells. If this is true, then increasing photoreceptor uptake of ketones may be a reasonable process to target for treatment in patients with unresolving MEKAR. If future animal studies confirm that drug-induced increases in RPE phagocytosis contribute to MEKAR, reducing phagocytosis of shed outer segments or decreasing production of ketones by RPE would not be recommended strategies for treatment as disruptions of both processes are associated with blinding conditions such as age-related macular degeneration (AMD), retinitis pigmentosa (especially those caused by mutations in *MERTK*), and diabetic retinopathy (16, 17).

### Selumetinib treatment decreases expression of AQP1 in hiPSC-derived RPE cells

Aquaporin 1 (AQP1) is a member of a family of transmembrane proteins that facilitate water transport. It has documented expression and functionality in the RPE derived from both human native tissue and differentiated from stem cells (18, 19). Because of its localization to the apical cell surface, it transports water to the basolateral direction, thus preventing subretinal fluid accumulation. Because of AQP1’s location and role in the RPE, it has been hypothesized that altered AQP1 may be responsible for causing MEKAR (5). Our results support this hypothesis with 10 μM of selumetinib causing decreased AQP1 protein expression following 10 days of selumetinib treatment. Bulk RNA-sequencing data revealed that *AQP1* gene expression was also significantly decreased in hiPSC-derived RPE treated with selumetinib (adjusted p=2.95x10^-4^).

Interestingly AQP1 levels normalized by 20 days of treatment, which parallels the clinical phenotype of patients who spontaneously resolve their MEK inhibitor-induced subretinal fluid accumulation. Our results contrasts with findings reported by Jiang *et al.* (20), in which they found that when ARPE-19 cells were treated with H2O2 or ultraviolet radiation, MEK activation led to a decrease in AQP1 protein levels. In turn treating cells with 1 μM of MEK inhibitor increased AQP1 expression. While both Jiang *et al.* and the present study implicate MEK activity in regulating AQP1 levels in RPE cells, the observed differences could be due to cell type (immortalized ARPE-19 cell line versus hiPSC-derived RPE) or pharmacological MEK inhibitor (preclinical research-grade MEK inhibitors PD98059 and U0126, versus clinically used selumetinib) (24). If future research *in vivo* demonstrates that decreased AQP1 contributes to MEKAR, a potential treatment for affected patients may be administration of agents that increase water transport via AQP1. For instance, drugs such as atrial naturietic peptide (ANP) or membrane-permeable analogs of cyclic GMP (cGMP), which Baetz *et al*. found significantly increase net fluid flux (25), may be useful for preventing MEKAR. Topical and oral administration of carbonic anhydrase inhibitors may be another potential therapeutic as its use decreases subretinal fluid in conditions such as central serous chorioretinopathy, retinitis pigmentosa, and X-linked retinoschisis (26–33). While the mechanism is unclear, it is thought to restore apical-to-basal fluid transport across RPE cells by acidifying the subretinal space (27, 29, 34, 35).

### Selumetinib decreases expression of genes regulating small molecule and ion transport as well as lysosomal acidification

The most striking difference in RNA-sequencing data between untreated and selumetinib-treated cells was the significant decrease in expression of genes involved in transport of ions and small molecules as well as those involved in lysosomal acidification (**Supplementary Table 1**). Depending upon the polar distribution of the affected genes’ protein products, the observed decrease in expression may be a direct effect of treatment with selumetinib or a cellular compensation (36).

*LRRC8C* codes for a subunit of the LRRC8 which is a volume-regulated anion channel (VRAC) (37). It contributes to cell volume homeostasis via its expression on the plasma membrane. When cells swell, this channel is activated and allows efflux of charged and uncharged molecules. However, regulatory volume decrease (RVD) is mainly mediated by efflux of potassium, chloride, and H2O. VRACs like LRRC8 primarily efflux chloride ions. LRRC8 has also been shown to be expressed on lysosomes, the “osmometer” of mammalian cells (38). LRRC8 contributes to lysosomal currents and is necessary to form large lysosome-derived vacuoles which store and eliminate water via exocytosis. In this way, they further maintain H2O balance intracellularly and have been described as the “bladder” of the cell. While the polarized localization of LRRC8 within the RPE cell has not been documented, decreased expression of *LRRC8C* in selumetinib treated hiPSC-derived RPE cells may prevent formation of these water-filled vacuoles, leading to apical fluid accumulation. Additionally, decreased expression of *LRRC8C* may be a cellular compensatory mechanism to inhibit efflux of chloride and water apically or the subretinal space *in vivo*.

*CLCN5* codes for ClC-5, which is a chloride-hydrogen exchanger with documented function acidifying lysosomes and up-taking protein in the proximal tubule cells (PTCs) of the kidney (39). Mutations in *CLCN5* cause Dent disease, a proximal tubulopathy of the kidney characterized by high levels of protein and calcium excreted in the urine, contributing to kidney stone formation. In PTCs, CIC-5 interacts with proteins important for receptor-mediated endocytosis and transportation to lysosomes for metabolism. Its expression on the cell membrane is thought to contribute to chloride currents. Polarized localization within RPE cells has not been documented. Decreased *CLCN5* expression in selumetinib-treated hiPSC-derived RPE could contribute to decreased lysosome acidification and may be a compensatory mechanism to prompt absorption of fluid by acidifying the subretinal space (35).

*SLC16A1* encodes MCT1, a H^+^-coupled monocarboxylate transporter that transports lactate into the cell (40). Due to its coupling with hydrogen ions, it contributes to intracellular acidification in addition to fluid transportation. MCT1 is highly expressed on the apical surface of RPE cells to facilitate uptake of metabolic waste products from overlying photoreceptors (40–42). Thus, decreased expression of MCT1 protein on the apical surface of RPE treated with selumetinib could contribute to decreased acidification intracellularly and accumulation of protons in the subretinal space. While protons attract water molecules, acidification of the subretinal space may prompt fluid transport across RPE cells and thus serve as a compensatory mechanism (35).

*SLC6A6* codes for a sodium- and chloride-dependent transporter (TauT) of taurine and beta-alanine localized to the plasma membrane (43, 44). Taurine is highly abundant in the retina, particularly in photoreceptors at the level of the outer nuclear layer. It is necessary for proper retinal structure and function with low levels associated with retinal degeneration at the level of the photoreceptors in animal models and humans (45–47). TauT is a well-known contributor to osmoregulation, whereby hypertonic extracellular conditions lead to intracellular influx of taurine and obligate water molecules mediated by transporters like TauT. TauT protein is expressed at similar levels in apical versus basolateral sides of human fetal RPE cells (48). Activity and gene and protein expression of TauT is increased in ARPE-19 cells following exposure to hyperosmolar media (46). Decreased expression of *SLC6A6* in selumetinib-treated cells may been a compensatory mechanism whereby cells with increased internalized POS decrease the amount of osmotically active taurine influx to prevent cell swelling. Thus, decreased permeability to taurine and water could lead to an accumulation of these molecules in the subretinal space.

*SLC22A23* encodes one of the ∼30 members in the highly conserved SLC22 transporter family (49). Its protein product’s function and structure have not been well-characterized. However, evolutionary analysis indicates that it likely functions as an organic anion transporter (OAT) and contributes to metabolic regulation. More work is needed to determine the effect of decreased *SLC22A23* observed in selumetinib-treated hiPSC-derived RPE cells.

*SLC12A2* encodes NKCC1, a Na^+^-K^+^-Cl^-^ cotransporter that mediates influx of chloride in an electrically neutral transfer, which contributes to cell volume regulation (50, 51). Its expression is localized to the basolateral membrane of chloride-secreting epithelial cells except in the olfactory epithelium, choroid plexus, and RPE, where is localized on the apical membrane (51–55). NKCC1 regulates subretinal fluid volume via ion transport (51). Because of its apical location, it transports chloride and therefore H2O out of the subretinal space in the basolateral direction. Thus, decreased expression of NKCC1 in selumetinib-treated hiPSC-derived RPE could decrease chloride and water transport from the apical surface and contribute to subretinal fluid accumulation.

*ATP6V1C2* encodes for a subunit of vacuolar ATPase (V-ATPase), a highly conserved proton pump which acidifies intracellular organelles, including lysosomes (56, 57). It hydrolyzes ATP to drive protons across a membrane. Specificity of localization in organelles and cell and tissue type are determined by isoforms of subunits. *ATP6V1C2* encodes for the V1C2 isoform. V-ATPases are found ubiquitously in the endomembranes of organelles and in the plasma membranes of certain cell types including osteoclasts, renal intercalated cells, male reproductive tract epithelial cells, and tumor cells. Multiple diseases are associated with mutations, mis-localization, and overexpression of V-ATPase subunits (57). Notably, a potentially pathogenic mutation was identified in *ATP6V1C2* (58). The missense mutation causing loss-of-function on V-ATPase was found with whole-exome sequencing in a patient with distal renal tubular acidosis. V-ATPases are found on the apical side of α-intercalated cells in the distal nephron and acidify urine. (57, 59). *ATP6V1C2* expression in RPE is documented (60–64). Animal models with inhibition or mutations of V-ATPases in the lysosomes of RPE cells impairs phagocytosis and degradation of photoreceptor outer segments (65–67).

### Selumetinib alters gene expression of aquaporins

Aquaporins are a family of membrane channels permeable to water. Treatment of hiPSC-derived RPE with selumetinib significantly altered gene expression of several aquaporins with confirmed expression in hiPSC-derived RPE: *AQP1, AQP3, AQP4, AQP7,* and *AQP11* (**Supplementary Table 2**) (19). Expression of *AQP1* and *AQP11* were increased, whereas expression of *AQP3, AQP4,* and *AQP7* decreased.

Interestingly, MAPK pathway activation helps cells cope with osmotic stress (68). Thus, inhibition of this pathway with selumetinib could potentially disrupt RPE cells’ ability to adapt to subretinal fluid accumulation by altering transcription of aquaporin genes. Notably, the ionic and osmotic composition of MEKAR-associated subretinal fluid has not been characterized. Regardless, alterations in expression of aquaporins in RPE cells due to MEK inhibition could certainly contribute to subretinal fluid accumulation.

While AQP3 protein expression has been confirmed in primary RPE cells, the polarized cellular localization has not been reported (69). Our results showed that MEK inhibition with selumetinib significantly increased *AQP3* expression in hiPSC-derived RPE. This contrasts with what Hollborn *et al*. found (69). They reported increased AQP3 mRNA induced by MEK activation in primary human RPE cells treated with CoCl2 to chemically induce hypoxia. To determine that this change in AQP3 expression was mediated by MEK activation, they inhibited MEK with the preclinical, research-use only molecule PD98059 (20 μM), whereas our study used selumetinib which is used in clinical practice to treat patients (24). Future work is needed to determine the cellular localization of AQP3 to the apical and/or basolateral side of RPE and if there are differences in the changes in AQP3 expression with MEK modulation in different types of RPE cell cultures (primary versus stem-cell derived) and with different MEK inhibitors.

Our study found AQP4 gene expression significantly increased in hiPSC-derived RPE treated with selumetinib. The polar cellular location of AQP4 in RPE has not been documented. Perhaps increased expression in selumetinib-treated cells is a compensatory mechanism to transport water out of the subretinal space into the choroid underlying the RPE. This differs from a study in ARPE-19 cells where inhibition of MAPK with 10 μM UO126 did not rescue decreased AQP4 protein expression in response to hyperosmolar conditions (70).

Like AQP3, AQP7 is also permeable to small solutes like glycerol and exhibited increased gene expression in hiPSC-derived RPE cells treated with selumetinib. AQP7 enables efflux of glycerol, thus increased AQP7 could allow for accumulation of glycerol and water in the subretinal space (71, 72).

Similar to *AQP1* gene expression in hiPSC-derived RPE treated with selumetinib, levels of AQP11 mRNA decreased. Research on Müller glia cells have demonstrated an association between intracellular edema and decreased AQP11 levels of activity contributing to Diabetic Macular Edema (DME) characterized by subretinal and intraretinal fluid accumulation (73). If AQP11 proteins behave the same way in RPE cells, then perhaps decreased expression in selumetinib-treated cells prevents water efflux. This increase in cell swelling may then prevent increased intracellular uptake of water, contributing to subretinal fluid accumulation.

### Comparison of MEKAR to ocular phenotypes caused by mutations in BEST1

*BEST1* encodes a membrane protein that acts as a cell volume-sensitive calcium-gated chloride channel which inhibits voltage-gated calcium channels (74, 75). While it is predominantly expressed in human RPE, it has also been detected in the kidney, brain, spinal cord, and testis (76, 77). Within the RPE, it is localized to the basolateral membrane and exhibits higher expression levels outside of the macula (78, 79). Interestingly, bestrophin protein is also suspected to regulate intracellular calcium and processes like phagocytosis and lysosomal function (80, 81). As demonstrated in this study, calcium signaling and regulation of cytosolic calcium concentrations were some of the top pathways affected in hiPSC-derived RPE cells. Pathologic mutations in *BEST1* can cause a spectrum of ocular phenotypes, mostly leading to detachment of the neurosensory retina at the macula with accumulation of subretinal fluid, causing a decrease in visual acuity (74). Notably however, this decrease in visual acuity is not large and many patients remain stable without treatment at 20/50. Like MEKAR, it is hypothesized that accumulation of fluid in the subretinal space in *BEST1*-associated diseases is due to dysfunction of RPE’s ability to transport ions and regulate fluid homeostasis including via phagocytosis of photoreceptor outer segments (POS). In contrast to MEKAR, it is thought that *BEST1* mutations modulating intracellular calcium levels may decrease uptake of POS, leading to accumulation of toxic lipofuscin in the subretinal space, killing both photoreceptors and RPE cells. Lipofuscin is a mixture of byproducts form POS degradation and includes oxidized proteins, lipids, and fluorophores. A study using hiPSC-derived RPE cells from patients with Best disease and non-affected siblings produced results that supported this hypothesis (82). They found cells from affected patients demonstrated decreased apical to basal fluid flux, increased accumulation of lipofuscin after feeding POS, decreased degradation of rhodopsin (RHO) after POS feeding, and differential responses to calcium stimulation. It is unknown whether lipofuscin accumulation is a feature of MEKAR. Given the clinical similarities between MEKAR and *Best1*-associated ocular diseases (eg, bilateral accumulation of subretinal fluid predominantly at the macula, decreased visual acuity, and affected proteins involved in ion transport, volume sensation and regulation, POS phagocytosis, and lysosomal acidification), future work should address the possibility of overlapping disease mechanisms.

## Conclusion

In summary, using an *in vitro* model of MEKAR, we have shown that the clinically used MEK inhibitor, selumetinib increases hiPSC-derived RPE cell phagocytosis, decreases AQP1 mRNA and protein, and alters gene expression with significant changes in genes involved in transport of ions and small molecules regulating cell volume and lysosomal acidification. Increased phagocytosis could potentially increase return of ketones to the subretinal space while decreased AQP1 protein expression would prevent water transport out of the subretinal space. This decrease is AQP1 may be a cellular response to prevent cell swelling of RPE accumulating internalized photoreceptor outer segments and their metabolites intracellularly. Thus, theoretically both increased phagocytosis and decreased AQP1 could contribute to the subretinal fluid accumulation characteristic of MEKAR. Future work is needed to further investigate these hypotheses and the roles of genes involved in lysosome acidification and cell volume regulation via molecular and ionic transport in the context of MEKAR.

### Future Directions

More work is needed to further assess whether the MEK inhibitor-induced increase in hiPSC-derived RPE cell phagocytosis affects the rate of ketone recycling into the subretinal space. This could be achieved using mass spectrometry on apical and basal cell culture media following the phagocytosis assays. This has been documented as the most reliable method to quantify secreted β-hydroxybutyrate (β-HB), the predominant ketone, from cultured human fetal RPE cells (83). Additionally, monocarboxylate transporter isoform 1 (MCT1) permits secretion of β-HB and is located on the apical side of RPE cells (84). Thus, quantification of MCT1 could also provide insight into an increased ability of ketone bodies to leave RPE cells and travel into the subretinal space. This would be especially interesting to quantify given that our RNA-sequencing data showed a significant decrease in *SLC16A*, the gene encoding MCT1.

This study assessed the effect of selumetinib on mRNA expression levels. There is precedent in the literature to evaluate changes in total RNA in future studies given the role of microRNAs (miR) in regulating phagocytosis (85). Specifically, miR-302d suppresses phagocytosis in cultured cells from the ARPE-19 cell line (86). Additionally, other miRs are reported to alter RPE autophagy (85). Thus, future work could quantify the impact of MEK inhibitors on microRNAs in cultured RPE cells.

Finally, the MEK inhibitor-induced differences in gene expression revealed in this study warrant further exploration. Specifically, work is needed to investigate if genes with altered expression play a role in the observed changes in hiPSC-derived RPE cell functions. For example, osmotic challenge assays and imaging can assess the effect of selumetinib on fluid flux and cell volume (18, 36).

## METHODS

### Sex As a Biological Variable

Cells from a single male donor were used in this study. There is no evidence to suggest that there is a sex difference in patients affected with MEKAR.

### Creation and Maintenance of Human Induced Pluripotent Stem Cell Cultures

Induced pluripotent stem cells were generated from a dermal biopsy of an unidentifiable male donor as previously described by Wiley *et al*. (14). All cell types were cultured at 37C, 5% CO2, and 20% O2. Briefly, fibroblasts were isolated and expanded from a 3 mm dermal biopsy and transduced with CytoTune-iPSC 2.0 Sendai Reprogramming Kit (Invitrogen/Thermo Fisher Scientific, Waltham, MA, USA). hiPSC colonies were isolated and expanded. TaqMan Human Pluripotent Stem Cell Scorecard Panel (Life Technologies/Thermo Fisher Scientific, Waltham, MA, USA) was used to analyze pluripotency and loss of viral transgene expression. Karyotype analysis was carried out by the University of Iowa Shivanand R. Patil Cytogenetic and Molecular Laboratory following a standard G-banding protocol.

### Differentiation and Maintenance of hiPSC-derived Retinal Pigment Epithelial Cells

We followed the rapid, directed differentiation protocol by Foltz and Clegg (15). The ROCK-inhibitor Y-27632 (MilliporeSigma, Burlington, MA, USA) was added to cell media at each passage for 4-7 days to increase cell survival and attachment. Briefly, hiPSCs were grown to 80% confluence in Essential 8 Flex media (Gibco/Thermo Fisher Scientific, Waltham, MA, USA) on rhLaminin-521 (Gibco/Thermo Fisher Scientific, Waltham, MA, USA) coated plates and then passaged with ReLeSR™ (STEMCELL Technologies, Vancouver, BC, Canada) 1:4 onto Matrigel-coated plates and treated for 14 days with growth factors. After RPE expansion, cells were maintained in RPE Supporting Media (RSM), composed of X-VIVO 10 (Lonza, Basal, Switzerland) and 0.2% Primocin (InvivoGen, San Deigo, CA, USA) on tissue culture-treated plates coated with Matrigel (60 ug/ml; Corning Life Science, Tewksbury, MA). Following the ROCK inhibition-dependent extended passage protocol by Croze, Bucholz, Radeke *et al*.(87), cells were passaged after reaching confluence every 4-5 days with TypLE™ Express (Thermo Fisher Scientific, Waltham, MA, USA) and plated at a seeding density of 2.5x10^4^ cells/cm^2^. Only cells at or below passage 5 were used to avoid epithelial-to-mesenchymal transition and loss of RPE morphology (e.g., pigment, cobblestone appearance) (87). Once cells were seeded onto experimental plates at density of 1.0x10^5^ cells/cm^2^ they were allowed to mature for 30 days before drug treatment, unless otherwise specified (e.g. TEER assay).

### Pharmacologic Inhibitor of the MAPK/ERK Pathway

Selumetinib (MEK inhibitor) was reconstituted to 10 mM in DMSO (Fisher Chemical, Hampton, NH, USA) per the manufacturer’s recommendations (Sigma-Aldrich, St. Louis, MO, USA).

### MTS Assay

To determine the maximum dose of selumetinib that could be applied to hiPSC-derived RPE cells without causing cell death, cells were plated in Matrigel-coated 96-well black/clear bottom plates (Corning Life Science, Tewksbury, MA) in X-VIVO 10. hiPSC-derived RPE were plated at 1.0x10^5^ cells/cm^2^ and allowed to mature for 30 days before starting drug treatment. Cells were treated with a log10-fold dilution from 1.0 x 10^-3^ – 1.0 x 10^3^ uM selumetinib for 3 days in media free of antibiotics. After 72 hours of treatment, selumetinib-free media was added to the cells along with 20 ul of CellTiter 96® AQqueous One Solution per well as directed in the manufacturer’s instructions (Promega, Madison, WI, USA). After 3 hours of incubation with MTS, the absorbance at 490 nm of each well was measured using the BioTek Cytation 5 microplate reader (Agilent Technologies, Santa Clara, CA, USA).

### Quantitative Real-Time Polymerase Chain Reaction (RT-qPCR)

cDNA was generated from 200 ng RNA using SuperScript IV VILO Master Mix (Thermo Fisher Scientific, Waltham, MA, USA). PCR reactions were run using PrimeTime™ qPCR probes (Integrated DNA Technologies, Coralville, Iowa, USA) and TaqMan™ Fast Advanced Master Mix (Thermo Fisher Scientific, Waltham, MA, USA). cDNA was amplified using a QuantStudio 6 Flex Real-Time PCR system (Thermo Fisher Scientific, Waltham, MA, USA). Data were analyzed using delta-delta Ct method in which values were normalized to the *B2M* housekeeping gene and corresponding mRNA values in hiPSCs from the same donor.

### Immunocytochemistry (ICC)

Cells were seeded onto 12-mm round glass coverslips coated in poly-L-lysine (Corning Life Science, Tewksbury, MA) and Matrigel which were placed in 24-well tissue culture treated plates (Corning Life Science, Tewksbury, MA). Cells were fixed with 4% paraformaldehyde for 20 minutes at room temperature then blocked for 1 hour. Cells were incubated with primary antibody for 2 hours at room temperature or overnight at 4C. Cells were washed with1xDPBS before incubating with secondary antibody and 1:2000 DAPI (Thermo Fisher Scientific, Waltham, MA, USA) for 1-2 hours at room temperature. Cells were washed then mounted onto microscope slides with coverslips and imaged at 40x using a Leica TCS SPE upright confocal microscope system (Leica Biosystems, Wetzlar, Germany). Primary and secondary antibody details shown in **Supplementary Table 3**.

### Transepithelial Electrical Resistance Assay (TEER)

Cells were seeded onto 6.5 mm cell culture inserts (Corning Life Science, Tewksbury, MA) coated with Matrigel. Five days after plating, cells were treated with 10 uM selumetinib for a total of 20 days. Resistance measures were obtained every 3 days using the EVOM Meter following the manufacturer’s protocol (World Precision Instruments, Sarasota, FL, USA). The resistance value of a cell culture insert without cells was subtracted from all sample resistance measurements.

### Rod Outer Segment Phagocytosis Assay

Bovine rod outer segments (bROS) (InVision BioResources, Seattle, WA, USA) at a concentration of 10 bROS/cell were added to hiPSC-derived RPE cells in a volume of 500 ul/well of a 24-well plate. Cells were incubated with bROS for 3 hours at 37C and 5% CO2. After 3 hours, all conditions were washed five times with 1xDPBS with calcium and magnesium. Prior to these washes, wells for the Internalized bROS condition were incubated with 2 mM EDTA in PBS for 10 minutes at 37C, 5% CO2 to remove unbound bROS. After washes, Total and Internalized condition wells were collected. 3 h and 24 h Chase conditions received fresh media with or without selumetinib per treatment group. At Chase collection time, cells were washed twice with 1xDPBS. Before collecting cells from any condition, 10 uM recombinant human FGF-basic (Peprotech/Thermo Fisher Scientific, Waltham, MA, USA) was added for a 10 minute incubation at 37C, 5% CO2. This step was done to ensure visualization of Ph-MAPK on Western Blot.

### Scanning Electron Microscopy

Cells were seeded onto 6.5 mm cell culture inserts (Corning Life Science, Tewksbury, MA). At Day 10 of treatment, cells were fed bROS and incubated for 3 hours at 37C and 5% CO2. Cells were fixed with 2.5% glutaraldehyde for 3-5 days then rinsed with 1xPBS. Samples were post fixed for 1 hour with 1% osmium tetroxide and rinsed with ddH2O. The samples underwent alcohol series dehydration followed by chemical drying with hexamethyldisilazane and left to dry in the fume hood for ≥ 2 days. Plates and membranes were mounted with colloidal silver onto standard SEM stubs and dried overnight. Sputter coating deposition was performed for 4 minutes with a current of 7mA. Coverslips were imaged on the Hitachi S-4800 at the Central Microscopy Research Facility.

### Transmission Electron Microscopy (TEM)

Cells were seeded onto 6.5 mm cell culture inserts with 0.4 um pore size (Corning Life Science, Tewksbury, MA). At Day 10 of treatment, cells were fixed overnight in ½ strength Karnovsky’s fixative. After primary fixation, samples were post fixed in osmium tetraoxide and uranyl acetate, and were dehydrated through a graded series of alcohols, acetone, and finally propylene oxide. The samples were then embedded in Spurr’s resin and cured at 60°C. After samples were fully cured ultrathin sections were collected onto copper square mesh grids using an ultramicrotome. Samples were imaged on the Hitachi HT7800 at the Central Microscopy Research Facility.

### Western Blot Analysis

Cells cultured and treated on 24-well plates were collected then lysed with 75 ul RIPA Lysis and Extraction Buffer with 1:100 Halt™ Phosphatase and Proteinase Cocktail Inhibitors (Thermo Fisher Scientific, Waltham, MA, USA). Protein concentration was quantified using Pierce™ BCA Protein Assay(Thermo Fisher Scientific, Waltham, MA, USA) with duplicates of 10 ul per standards and samples. Colorimetric changes in plates were read at 562 nm with the BioTek Cytation 5 microplate reader after 30 minutes of incubation at 37C. 10 ug protein was run on a 4-20% Novex™ Tris-Glycine gel (Thermo Fisher Scientific, Waltham, MA, USA) for 25 minutes at 225 volts. Broad-range protein transfer to blots was performed using iBlot™ 3 set to a 6 minute run at 25 volts with low cooling (Thermo Fisher Scientific, Waltham, MA, USA). After incubation with blocking buffer for 1 hour at room temperature or overnight at 4C, blots were probed with primary antibody then washed and probed with HRP-conjugated secondary antibody (as shown in **Supplementary Table 4**). Blots were imaged using the iBright^TM^

FL1500 (Thermo Fisher Scientific, Waltham, MA, USA). Blots were then stripped with Restore^TM^ stripping buffer (Thermo Fisher Scientific, Waltham, MA, USA) and probed for beta-actin after imaging to confirm stripping removed previously bound antibody for protein of interest. Densitometry was performed in FIJI ImageJ. All protein bands were normalized to beta-actin.

### Gene Expression Profiling Using RNA-Sequencing

RNA from cells was isolated using the NucleoSpin RNA kit (Machery-Nagel, Düren, Germany) and final samples were eluted in 40 ul of RNAse free water. Gene expression profiling using RNA-Seq was performed by the University of Iowa Genomics Division using manufacturer recommended protocols. Briefly, 250 ng of DNase I-treated total RNA was used to prepare sequencing libraries using the Illumina stranded mRNA library preparation kit (Illumina, Inc., San Diego, CA, USA).

The pooled and barcoded libraries were sequenced on the Element AVITI24 sequencing platform (Element Biosciences, San Diego, CA) using a Cloudbreak FS 150 cycle (2x75 bp) high output sequencing flow cell to generate at least 30M paired-end sequencing reads per sample.

### Bulk RNA Sequencing Analysis

Data were processed using the nf-core/rnaseq pipeline v3.18.0; doi: 10.5281/zenodo.1400710 from the nf-core workflow collection (88), utilizing reproducible software environments from the Bioconda (89) and Biocontainers (90) projects. The pipeline was executed with Nextflow v24.10.5 (91). In brief, paired-end sequence reads were pre-processed by Trim Galore v0.6.10 (92)using Cutadapt v4.9 (93) to remove adapter sequences and filter low quality reads. Both raw and trimmed FASTQ files (94) were evaluated for quality control using FastQC v0.12.1 (95) to assess Phred scores, GC content, sequence length distributions, and identify over-represented sequences. Lightweight pseudo-alignment and quantification was completed using Salmon v1.10.3 (96) with the selective alignment mapping strategy (97). A transcriptome index was constructed from Ensembl *Homo sapiens* primary assembly GRCh38 by generating a decoy-aware transcriptome for k-mers of length 31 from gene transfer format (GTF) genome annotations (release 113) and the full reference genome sequence. Paired-end reads were then quantified in mapping-based mode for inward reverse strand-specific libraries.

Quantification results were analyzed using R ((98); R version 4.4.1 (2024-06-14)) and several Bioconductor packages v3.20.0 (99). Transcript-level quantification results output by Salmon were summarized to gene-level abundance estimates and read counts (100) using tximeta v1.24.0 (101) with a linked transcriptome to the Ensembl GRCh38 reference genome. Additional gene annotations were obtained using AnnotationDbi v1.68.0 (102) and org.Hs.eg.db v3.20.0 (103). Exploratory data analysis was facilitated by data structures and methods from the SummarizedExperiment v1.36.0 (104) and tidybulk v1.18.0 (105) packages. Analysis of differential expression was limited to samples from day 10, 20, and 30 time points. Low count genes were excluded following (106), setting a minimum count-per-million threshold of 10 for a minimum group size of 3. Fold change estimates and p-values were calculated using DESeq2 v1.46.0 (107) by fitting a negative binomial generalized linear model with a Wald test to assess coefficient significance. To identify treatment-driven gene expression changes, an additive model was specified to account for batch (1, 2, or 3) and treatment (10 µM vs. 0 µM) effects (108). Pathway analysis was conducted with iPathwayGuide using BH-adjusted *p*-values with differential expression thresholds of 0.05 for genes with absolute log2 fold changes > 1 (109).

### Statistics

Statistical analysis was performed in GraphPad Prism Version 10.2.3 (347) unless otherwise noted. All data were collected from three independent experiments (n=3). Statistical significance was calculated using paired t-test for all RPE cell function assays (alpha = 0.05, FDR = 0.01).

### Study Approval

Cell obtained from an anonymous donor, exempt from human subjects approval.

### Data Availability

Data is available on Gene Expression Omnibus (GEO): accession number GSE308550.

## AUTHOR CONTRIBUTIONS

LPL, RFM, EMB, and BAT conceptualized the study, devised methodology, and generated figures. LPL conducted experiments, performed formal analysis, and wrote the original draft. JAR performed formal analysis, generated figures, and submitted data to Genome Expression Omnibus. LPL, MJ, RJ, LAW, BH, MJR, and ATW acquired data. RFM, EMB, BAT, BAE, HCB and TMB reviewed and edited the manuscript. RFM, EMB, and BAT provided resources and supervised the study. EMB and BAT acquired funding.

## ACKNOWLEDGEMENTS

The authors thank the patient for their tissue donation and generosity in supporting biomedical research. This work was funded by the Gilbert Family Foundation, P30 EY025580, and the University of Iowa Medical Scientist Training Program (T32 GM139776). Data presented herein were obtained at the Genomics and Bioinformatics Divisions of the Iowa Institute of Human Genetics (RRID: SCR_023422) which is supported, in part, by the University of Iowa Carver College of Medicine. We would like to thank the directors of each of these divisions for their guidance, Kevin Knudtson and Michael Chimenti, respectively. We offer particular thanks to Mary Boes who performed bulk RNA-sequencing. We also would like to thank the Central Microscopy Research Facility for providing instrumentation for scanning and transmission electron microscopy. Figures generated with BioRender.

