## Supplementary Data for "Modeling MEK inhibitor-Associated Retinopathy *in vitro* using human induced pluripotent stem cell-derived retinal pigment epithelial cells"

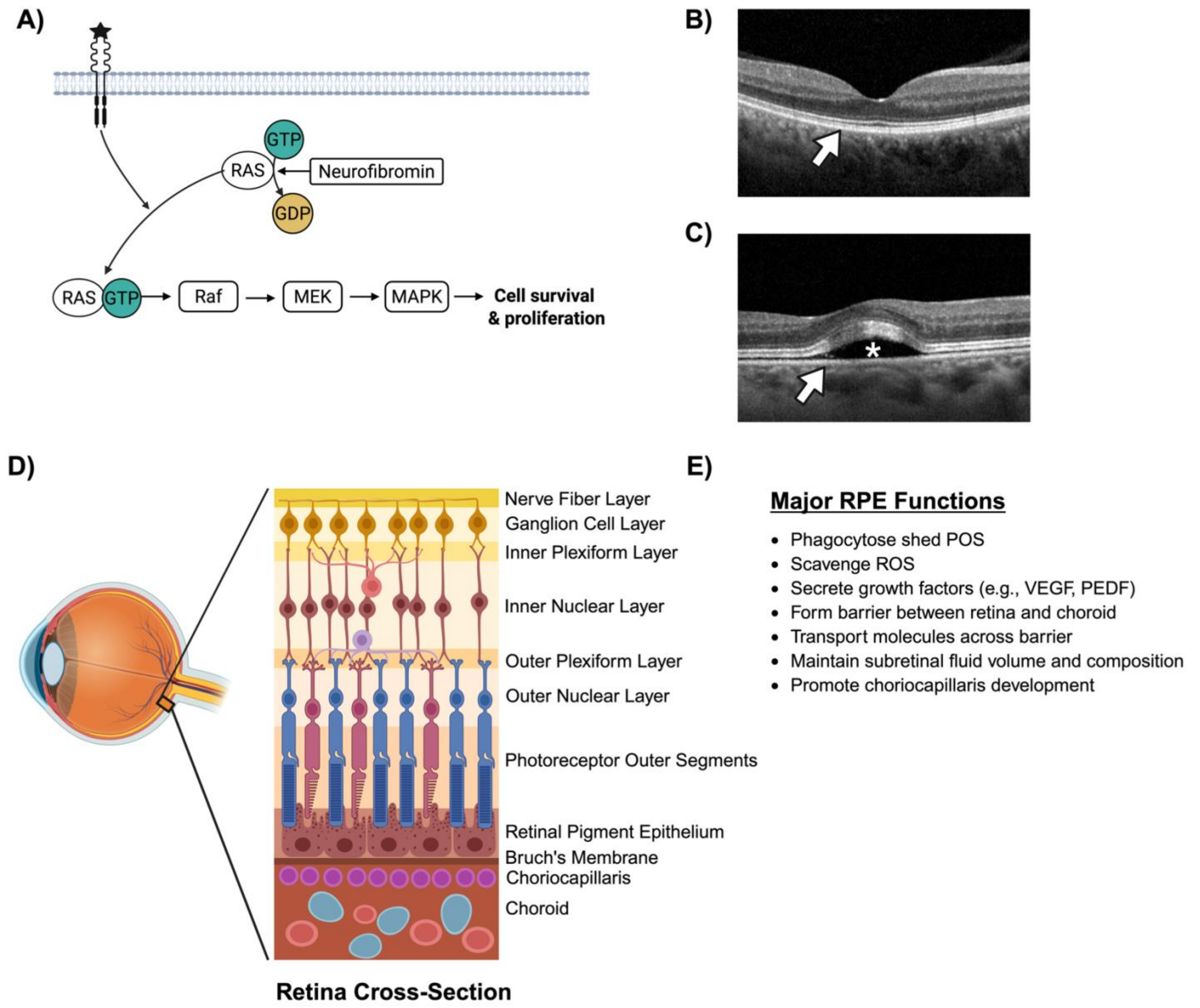

**Supplementary Figure 1 – Overview of MEK inhibitor-Associated Retinopathy** A) MEK inhibitors are anti-cancer drugs designed to suppress activity of the MAPK pathway. B) OCT of normal retina and retinal pigment epithelium (RPE) (arrow). C) Drug-induced retinopathy often occurs in both eyes at the fovea (responsible for central vision) within the first two weeks of starting treatment and is characterized by accumulation of fluid (asterisks) separating the retina from the underlying RPE (arrow). D) Cross-section of the posterior fundus depicting the retina, RPE, and choroid. E) Major functions of the RPE. Abbreviations: OCT, ocular coherence tomography; POS, photoreceptor outer segments; ROS, reactive oxygen species; VEGF, vascular endothelial growth factor; PEDF, pigment epithelium-derived factor.

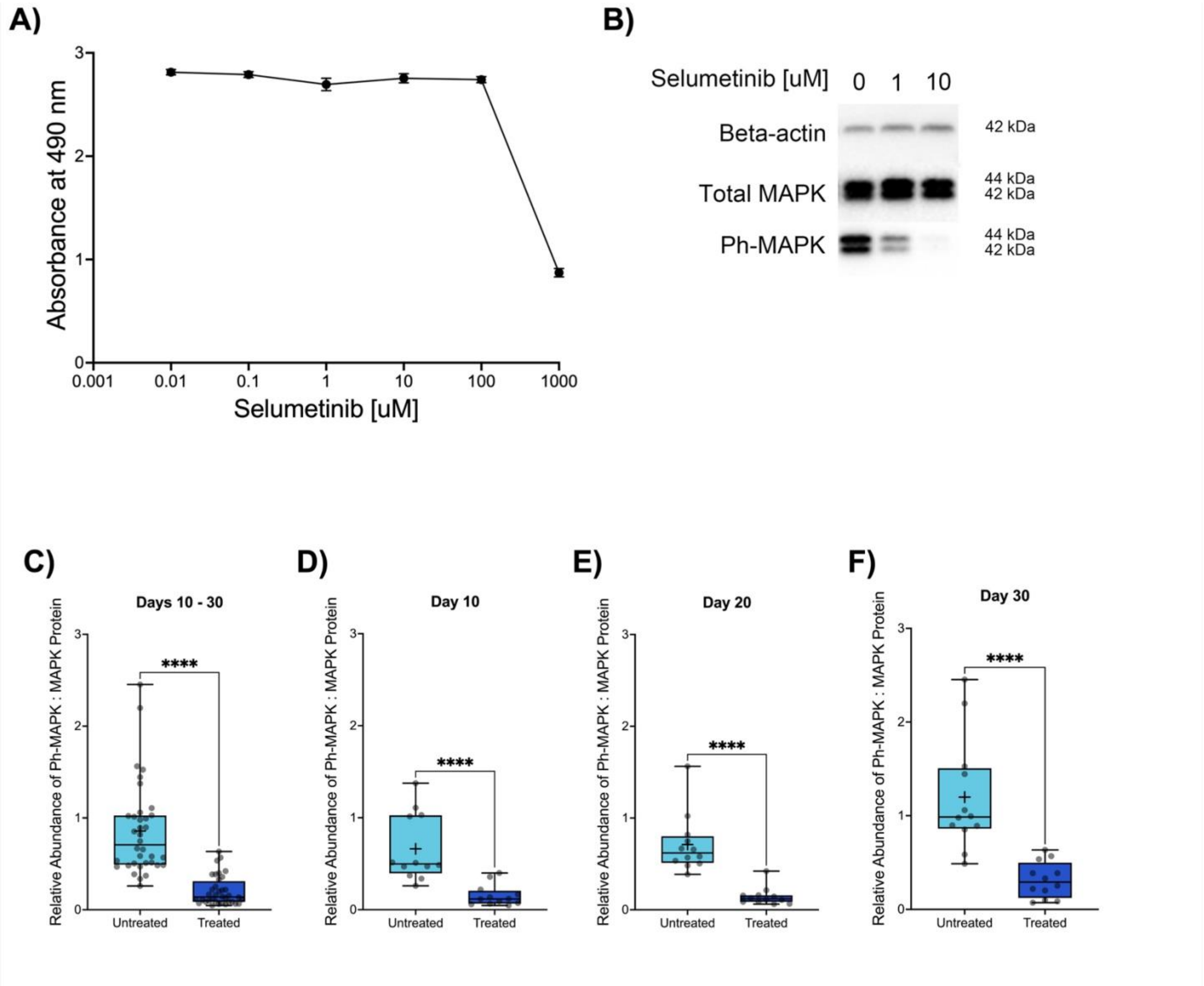

**Supplementary Figure 2 - Optimization of Selumetinib Dosage** A) Cell viability of hiPSC-derived RPE drops precipitously at doses greater than 100 μM selumetinib as indicated by decreased absorbance at 490 nm, representing inability of dead cells to metabolize MTS reagent. Mean ± SEM of 8 technical replicates. B) 10 uM selumetinib has greater inhibition of MAPK activation via phosphorylation compared to 1 uM selumetinib. C-F) Western blot densitometry indicates robust inhibition of MAPK phosphorylation cumulatively and when separated by collection time point. Paired t tests showing mean ± SEM, n = 36 pairs for (C), n = 12 pairs for (D-F). ****, $p<0.001$.

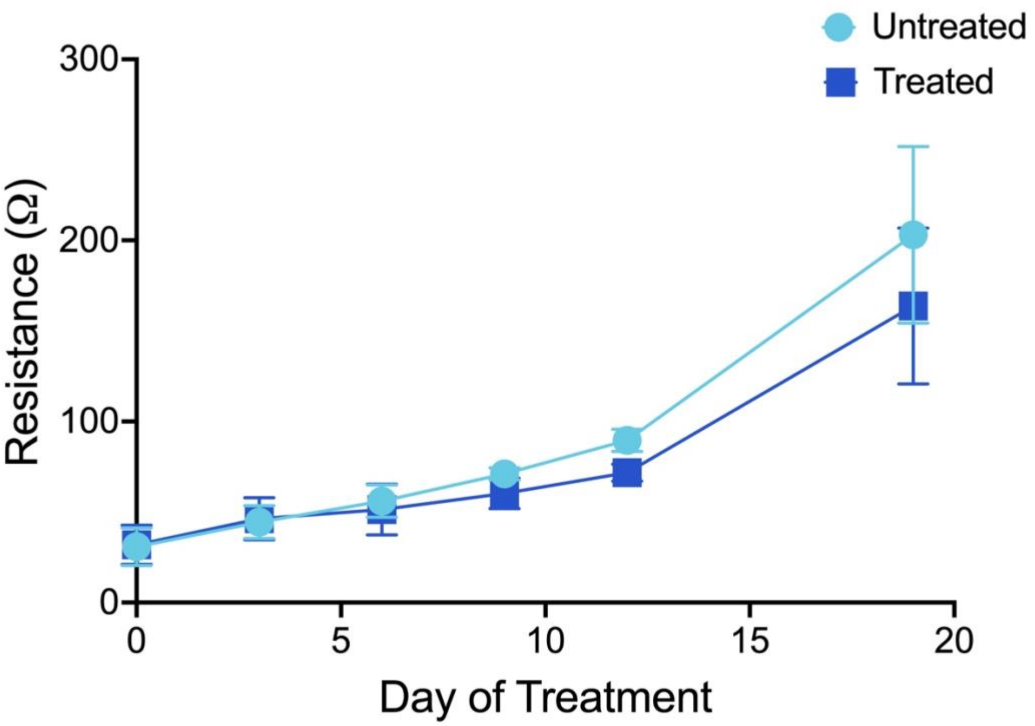

**Supplementary Figure 3 - Selumetinib does not affect TEER of hiPSC-RPE** Paired t test did not detect a difference between untreated and treated hiPSC-derived RPE at any time point. Mean ± SEM, n = 3 independent experiments.

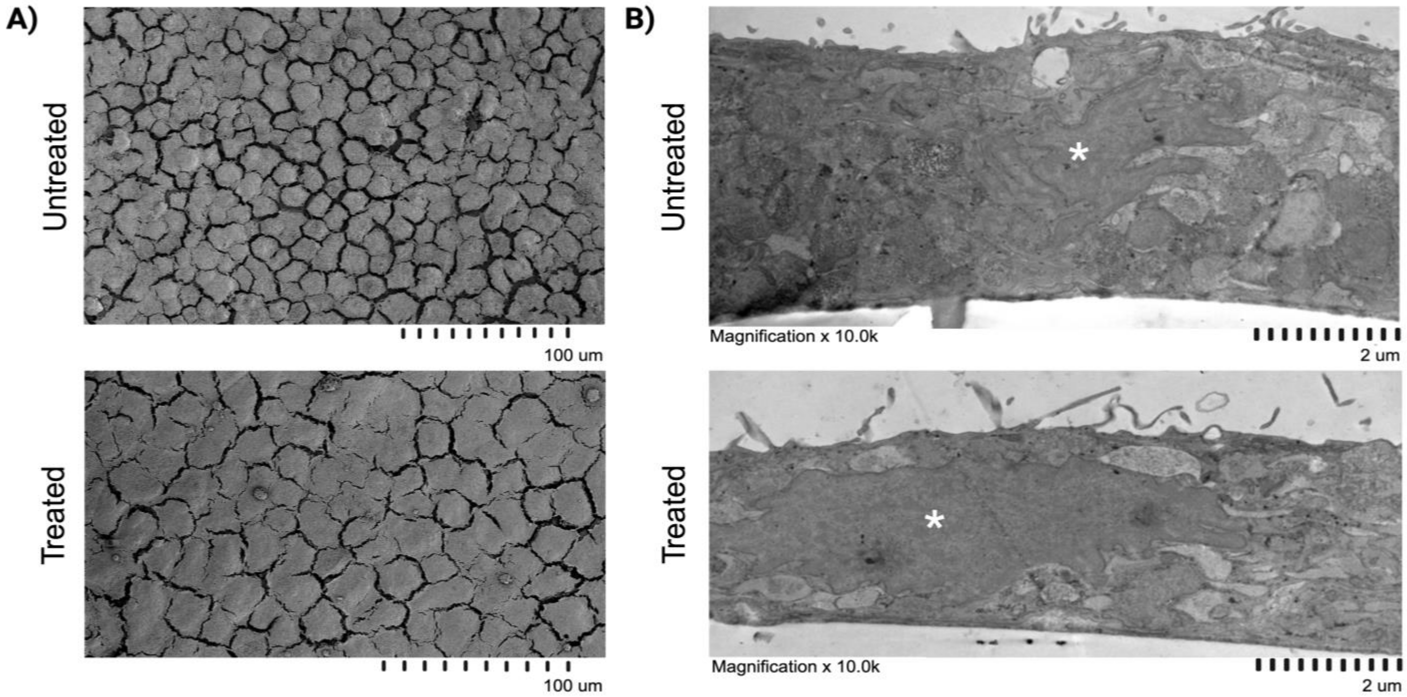

**Supplementary Figure 4 – Scanning and transmission electron microscopy of hiPSC-derived RPE. A)** Scanning electron microscopy of hiPSC-derived RPE cells. **B)** Transmission electron microscopy of hiPSC-derived RPE cells (nuclei indicated with asterisks).

| Rank in Top 50 DEGs | Gene | $Log_2$Fold Δ | Adjusted p-value | NCBI Gene Summary | Expressed in Human RPE? |
|---|---|---|---|---|---|
| 7 | LRRC8C | -1.25 | $2.00 \times 10^{-97}$ | "Enables volume-sensitive anion channel activity. Involved in cyclic-GMP-AMP transmembrane import across plasma membrane and monoatomic anion transmembrane transport. Located in cytoplasm and plasma membrane." (1) | Yes |
| 17 | CLCN5 | -1.01 | $5.59 \times 10^{-76}$ | "Member of the ClC family of chloride ion channels and ion transporters. The encoded protein is primarily localized to endosomal membranes and may function to facilitate albumin uptake by the renal proximal tubule." (2) | Yes |
| 23 | SLC16A1 | -1.32 | $1.21 \times 10^{-70}$ | "Proton-linked monocarboxylate transporter that catalyzes the movement of many monocarboxylates, such as | Yes |

| | | | | | |
|---|---|---|---|---|---|
| | | | | lactate and pyruvate, across the plasma membrane.” (3) | |
| 25 | SLC6A6 | -1.25 | $1.22 \times 10^{-69}$ | “A multi-pass membrane protein that is a member of a family of sodium and chloride-ion dependent transporters. The encoded protein transports taurine and beta-alanine.” (4) | Yes |
| 34 | SLC22A23 | -1.08 | $1.43 \times 10^{-63}$ | “Belongs to a large family of transmembrane proteins that function as uniporters, symporters, and antiporters to transport organic ions across cell membranes.” (5) | Yes |
| 42 | SLC12A2 | -1.66 | $9.15 \times 10^{-60}$ | “Mediates sodium and chloride transport and reabsorption. The encoded protein is a membrane protein and is important in maintaining proper ionic balance and cell volume.” (6) | Yes |
| 49 | ATP6V1C2 | -2.28 | $4.65 \times 10^{-58}$ | “A component of vacuolar ATPase (V-ATPase), a multisubunit enzyme that mediates acidification of | Yes |

| | | | | |
|---|---|---|---|---|
| | | | | eukaryotic intracellular organelles. V-ATPase dependent organelle acidification is necessary for such intracellular processes as protein sorting, zymogen activation, receptor-mediated endocytosis, and synaptic vesicle proton gradient generation." (7) |

**Supplementary Table 1 – Notable differentially expressed genes in hiPSC-derived RPE treated with selumetinib.** Genes that may contribute to observed clinical and experimental phenotype from the list of top 50 genes with significantly different expression between untreated and treated cells. Unless otherwise cited, expression in human RPE confirmed using IOWA integrated retina, RPE, and choroid (from studies between 2019 – 2020) dataset from Spectacle (8-12). Abbreviations: DEG, differentially expressed genes.

| Gene | $Log_2 Fold\ \Delta$ | Adjusted p-value | NCBI Gene Summary | Expressed in Human RPE? |
|---|---|---|---|---|
| AQP3 | 1.64 | $3.24 \times 10^{-50}$ | "Localized at the basal lateral membranes of collecting duct cells in the kidney. In addition to its water channel function, aquaporin 3 has been found to facilitate the transport of nonionic small solutes such as urea and glycerol, but to a smaller degree. It has been suggested that water channels can be functionally heterogeneous and possess water and solute permeation mechanisms." (13) | Yes (14) |
| AQP7 | 1.86 | $2.92 \times 10^{-14}$ | "Localized to the plasma membrane and allows movement of water, glycerol and urea across cell membranes. This gene is highly expressed in the adipose tissue where the encoded protein facilitates efflux of glycerol. In the proximal straight tubules of kidney, the | Yes (14) |

| | | | | |
|---|---|---|---|---|
| | | | encoded protein is localized to the apical membrane and prevents excretion of glycerol into urine. The encoded protein is present in spermatids, as well as in the testicular and epididymal spermatozoa suggesting an important role in late spermatogenesis." (15) | |
| AQP4 | 1.82 | $3.23 \times 10^{-7}$ | "Function(s) as water-selective channels in the plasma membranes of many cells…the predominant aquaporin found in brain and has an important role in brain water homeostasis." (16) | Yes (14) |
| AQP1 | -0.39 | $2.95 \times 10^{-4}$ | "(A) small integral membrane protein with six bilayer spanning domains…permits passive transport of water along an osmotic gradient. This gene is a possible candidate for disorders involving imbalance in ocular fluid movement." (17) | Yes (14) |
| AQP11 | -0.18 | 0.015 | "Involved in several processes, including glycerol | Yes (14) |

| | | | |
|---|---|---|---|
| | | | transmembrane transport; hydrogen peroxide transmembrane transport; and intracellular water homeostasis. Located in cell surface; endoplasmic reticulum; and plasma membrane.” (18) |

**Supplementary Table 2 – Differentially expressed aquaporin genes in hiPSC-derived RPE treated with selumetinib.**
